# Using a citizen science approach for research-informed teaching of evolution and palaeontology

**DOI:** 10.1101/2024.11.01.621579

**Authors:** Stephan Lautenschlager

## Abstract

Over the past few decades, citizen science has proven to be a valuable tool for research projects by involving the public in large-scale data collection. This collaboration between researchers and volunteers has been shown to be highly beneficial, allowing for efficient data collection in shorter time frames than possible for individual researchers. This study introduces a citizen science-inspired approach to teaching and curriculum design, involving undergraduate students directly in active research. Using a case study on dinosaur eye size, integrated into a first-year undergraduate course in Geology and Palaeontology at a UK university, the study presents the advantages of this approach for both teachers and learners. As part of the study, 22 undergraduate students were involved in data collection, analysis, and the subsequent publication processes, emphasizing active student participation in research. A subsequent survey demonstrated high student engagement and perceived relevance of the citizen science-style teaching approach. Results indicate that students found the approach appealing, engaging, and beneficial for understanding scientific concepts and methods. The case study shows that a citizen science approach to research-informed teaching can enhance student engagement and learning by providing meaningful, hands-on research experiences. This approach allows students to apply theoretical knowledge in a realistic context, fostering independence in quantitative research skills and collaborative learning. Despite potential challenges related to data accuracy and student motivation, the benefits of integrating such approaches into higher education curricula are substantial, offering a valuable model for teaching in Earth Sciences and beyond.

## Introduction

Over the last 60 years, there has been a significant evolution in the role of research and its intersection with teaching in higher education (Committee on Higher Education 1963; Elton, 1992). In particular, in the UK and other English-speaking countries, the paradigm of *research-led teaching* has emerged as the benchmark for bringing research into university lecture rooms (DfES, 2003; Schapper & Mayson, 2008), championed by research-intensive institutions aimed at securing a competitive edge in student recruitment (Healey et al., 2003). However, a clear definition of what constitutes research-led teaching remains more elusive, with various synonymous terminologies being used. In fact, numerous studies have been dedicated to expanding on the original concept and on the different forms of integrating research into teaching curricula (Healey et al., 2003; Griffiths, 2004): (I) *Research-led teaching involves* specialists educators (i.e. researchers) introducing research findings in taught modules, often selected from their area of expertise. (II) *Research-oriented teaching* places greater emphasis on the processes of how research knowledge is acquired and produced, often highlighting inquiry-based skills, and drawing from the experience of researchers or conventions and standards in the discipline. (III) *Research-based teaching* is centered around inquiry-based activities to achieve the learning objectives. (IV) *Research-informed teaching* brings all of these elements together with teachers and students actively engaging in inquiry-based activities underpinned by the teacher’s research background.

The first two definitions and approaches predominantly position students as passive audiences, receiving information they may find useful but cannot necessarily connect with. In fact, students may perceive research-led teaching as a disadvantage if they feel sidelined by the prioritisation of educator’s research activities over their own learning (Neumann, 1994; Jenkins et al., 1998; Zamorski, 2002; Healey et al., 2003). In contrast, research-based and research-informed teaching includes students as active participants in both their learning as well as research activities. A number of studies and papers have been dedicated to engaging students in learning and research going beyond the classic transmission model of just including and presenting research in the curriculum. For example, Brew (2002) makes a distinction between teacher-focused and student-focused learning, similarly arguing that the latter is beneficial with students being able to take ownership of their own learning. However, the question remains how students can be confronted with research content effectively and be motivated to engage with it. The approach presented in this work takes inspiration from a different example where researchers and learners (in the widest sense) engage and interact on a mutually beneficial basis - citizen science.

Broadly defined as involving the public in large-scale data collection efforts for specific research projects, citizen science has gained traction in recent decades. This is not a new concept, and although the term citizen science was coined about 30 years ago, some initiatives to have the public participate in scientific data collection date back to the 19th century; for example, the National Audubon Society’s Christmas Bird Count project has benefitted from thousands of volunteers collecting data on bird sightings for more than a century (Bonney et al., 2009). However, it is only in the last decades that citizen science has become a widespread tool, underpinned by rigorous scientific protocols and data collection metrics (Cohn, 2008). Employed mostly within natural and life science disciplines, numerous examples have now solidified citizen science as an important research tool to integrate the public and to maximise data collection.

Many research questions within biological and Earth sciences can have large-scale spatial, temporal, and taxonomic dimensions with study objects distributed globally or across geological timescales. Harnessing support from the public is therefore a clear advantage of citizen science, allowing data collection efforts beyond the capabilities of individual researchers. However, citizen science has not been free from criticism. Despite its advantages, citizen science has faced criticism including concerns regarding its potential for neoliberalisation, exploitation of unpaid labour, and commercialisation of the collected data (Michelucci & Dickinson, 2016; Vohland et al., 2019, 2021). Additionally, the top-down/scientist-driven approach of citizen science projects, where participants are relegated to mere data gatherers poses challenges to meaningful engagement (Powell & Collin, 2009). However, for large-scale projects, it may be logistically impossible to include participants in study design, data analysis, and dissemination.

In a classroom context, many of these problems can be avoided (or actually don’t exist in the first place). In contrast to normal citizen science projects, university students are not amateurs and already possess or are being taught the skills that could be relevant to research projects. I here present a citizen science approach to research-informed teaching exemplified by a recent case study. It is argued here that having students actively participate in research alongside teachers improves students’ understanding and knowledge of the subject and has a positive impact on student engagement and learning.

### Case study: Eye size of dinosaurs

#### Study design

The presented case study was integrated into a year 1 undergraduate course (“Geocience Project”) introducing and cultivating foundational skills in research methods (e.g. data collection, analysis, and presentation) as part of a Geology and Palaeontology degree at a research-intensive UK university. The course spanned an entire teaching semester of eleven weeks comprising 2-hour practical sessions each week, accompanied by lectures and corresponding practicals on statistics and data analysis for the first three weeks. The practical components focussed primarily on data collection and analysis for a research project after an introduction to the topic and the research methods to be used were provided in the first two weeks. For the academic year of 2022/2023, the chosen topic focussed on collecting data on the orbit (=eye socket) size of dinosaurs and related groups and on estimating eye size to quantify visual capabilities (Lautenschlager et al., 2023). The topic was selected based on the following: (i) alignment with the instructor’s research focus and expertise; (ii) availability of relevant measurements and data from the scientific literature; (iii) novelty of the research question, filling a considerable knowledge gap; (iv) an extensive dataset allowing for data collection by over 20 undergraduate students without unnecessary redundancy or overlap; (v) flexibility to accommodate students’ learning styles.

#### Data collection

Required data included measuring the length of the orbit and the skull length, alongside obtaining information on the taxonomic relationship, the time range, and the diet of each fossil species from literature sources (fig. 1). Each student was assigned a group of 20-30 species of the same taxonomic group (e.g. ceratopsian dinosaurs, theropod dinosaurs, pterosaurs) to get them started. Each group was assigned twice so that the accuracy of the data could be compared afterward, and outliers identified. All data was entered on a shared Google spreadsheet accessible to the entire student group. This data subsequently formed the basis for a publication with all students included as co-authors (Lautenschlager et al., 2023).

**Fig. 1.**
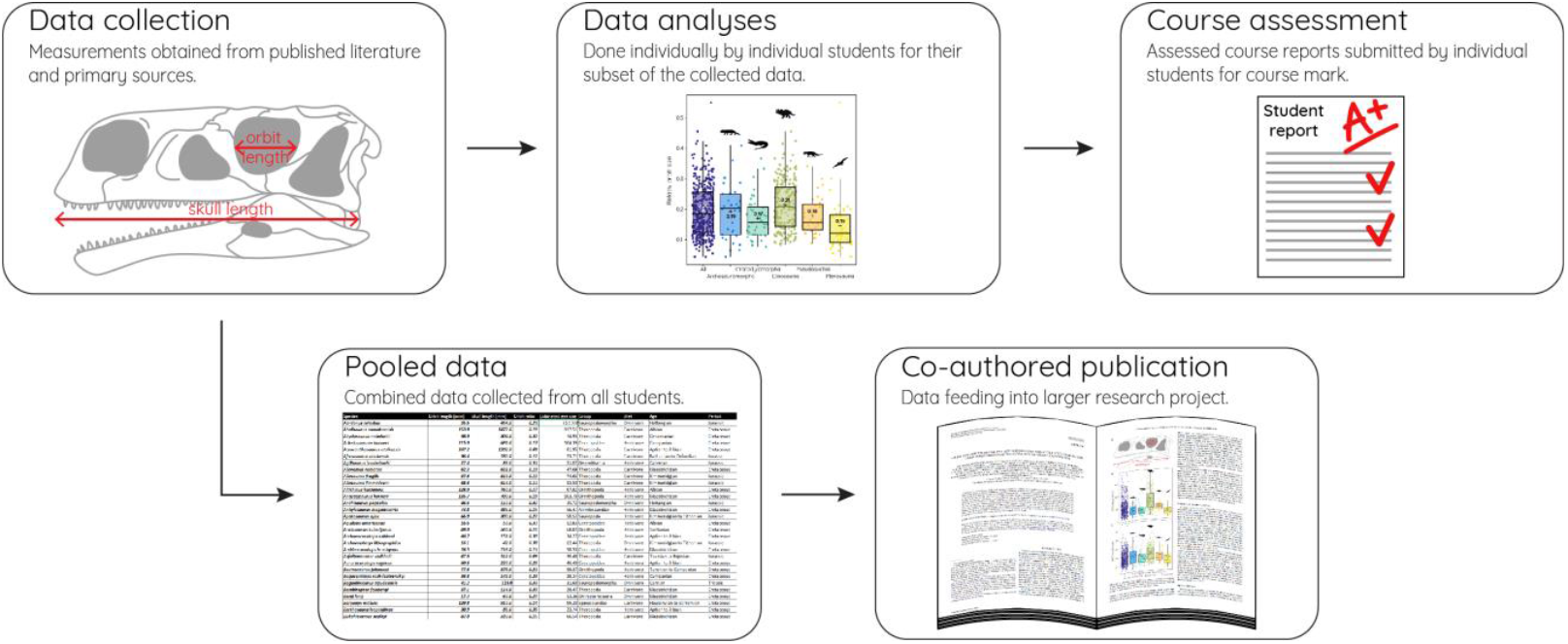
Schematic overview of the presented citizen science-style teaching approach.

#### Assessment

As a formal requirement for the degree programme, the course was assessed in the form of a 1500-word research report and a presentation. Students were given the opportunity to submit a draft version of their reports which was not marked and only received formative feedback. Students were given flexibility in selecting a topic for both their reports and presentations, provided the collected data was integrated in some form. Similarly, there was flexibility to choose appropriate statistical tests and to select plots and figures to present the results. Training in relevant software (Excel, PAST, R) was provided as part of the practical components. Students were allowed to compare their data to the entire data set or to subsets collected by their peers as part of the assessment. This approach maximised the flexibility of the assessed work, fostering a collaborative environment for students to leverage collective insight and data.

#### Learning goals

The overarching aim of the presented approach was to create a learning scenario with students fully emerged into a research project with applicability beyond the lifetime of the course. This was achieved by individual learning goals:

- Conducting data collection within a realistic research setting
- Applying appropriate statistical tests and data analyses
- Communicating research findings effectively through written, graphical, and verbal format
- Cultivating independence in quantitative research skills
- Maximising learning and engagement with the subject

#### Evaluation

In order to gain insight into how students perceived the citizen science-style approach they were sent a link to an anonymous online questionnaire. Students were asked to respond on a 5-level Likert-style scale (1 = no, absolutely not, 2 =no, 3 = neutral/indifferent, 4 = yes, 5 = yes, very much) to the following statements:

i. Data collected by students as part of the course was used for a research project and subsequent publication. Did this fact contribute to the course’s/activity’s appeal?
ii. Did the data collection activity improve your engagement with the course?
iii. Did the data collection activity foster your willingness to attend the course in person?
iv. Did the data collection activity as part of a larger research project help you with understanding scientific concepts and methods?
v. Did you find the data collection activity meaningful and relevant to your degree and career goals?
vi. Would you recommend that the activity is repeated each year in this course (with different data and research questions)?

## Results

### Data collection

At the end of the 11-week course, students collected data for over 350 different species of fossil archosaurs. Upon checking for correctness only a few measurements (for 27 species) had to be repeated as a comparison between the two collected sets of data showed some discrepancies. A few additional specimens were added to the data set as students had not taken measurements due to not being able to find the respective publications, lack of scaling information for the specimens, and difficulties with taxonomic assignments. The final data set contained 382 specimens and formed the basis for a publication. A manuscript was submitted in July 2023 (approximately three months after the course had ended) with students included as co-authors and involved in manuscript generation. A manuscript was published in January 2024 following two rounds of minor revisions (Lautenschlager et al., 2023).

### Student survey

The online questionnaire was completed by 16 out of 22 students representing a response rate of 72%. The 16 students responded to all six statements and responses are summarised in figure 2. All students agreed (5 students) or strongly agreed (11 students) that the data collection approach contributed to the course’s appeal. Similarly, the majority of students agreed/strongly agreed that the activity improved their engagement with the course and positively impacted their willingness to attend. Students further agreed (8 students) or strongly agreed (8 students) that the data collection helped them with understanding scientific concepts and methods. With regard to the relevance of the data collection approach for their degree and career goals, students were a bit more ambivalent (6 students strongly agreed, 6 agreed, 2 were indifferent) but the vast majority agreed that the activity should be repeated each year.

**Fig. 2.**
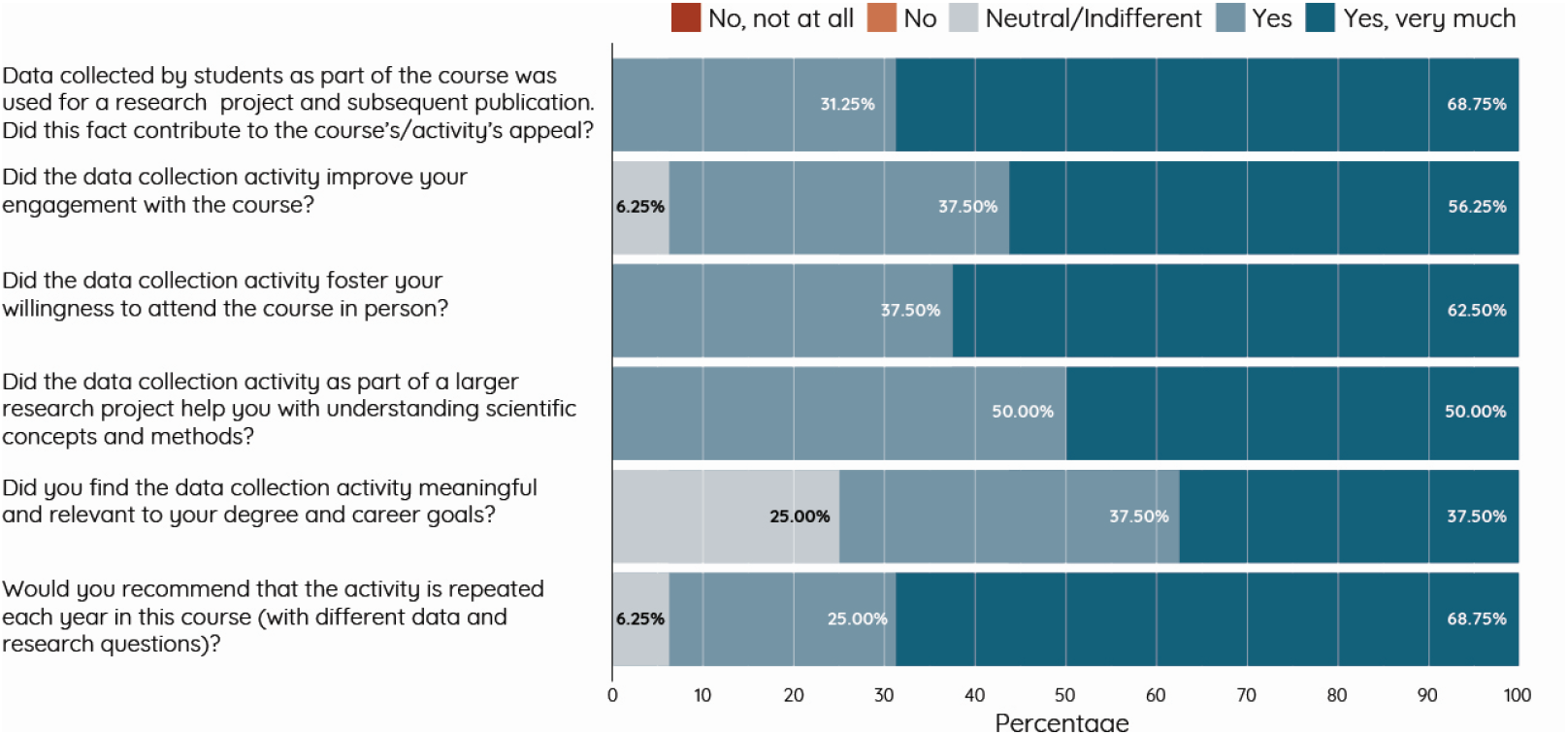
Summary of student responses to the survey questions about the conducted citizen science-style project.

## Discussion

### Benefits and disadvantages of a citizen science teaching approach

The citizen-science approach to teaching introduced here has several clear advantages. Survey responses demonstrate that students found the activity highly beneficial, particularly for understanding scientific concepts and methods; but also in terms of course engagement students rated the approach overwhelmingly positively. This aligns with research showing that students generally obtain a deeper understanding when applying their learning to real-world situations and tasks (Kuh, 2009). However, applicable and realistic activities do not automatically guarantee improved engagement; they need to be underpinned by explanations of the underlying principles and the introduction of core concepts (Collins, 1991; Carroll et al., 2008). While it may be possible to achieve this using mock data and/or examples, a citizen science approach provides a straightforward rationale by involving students in collaborative data collection as part of a larger research project.

Additionally, the citizen science approach allows for accommodating different forms of student learning. Students were able to work independently or collaboratively discussing best approaches and comparing collected data

The experience gained from this activity can further be beneficial in a longer-term context, particularly for dissertation projects and possibly future careers. However, while student responses were generally positive, some were uncertain about the activity’s relevance to their career goals. This uncertainty may stem from several reasons: The case study was designed within the context of academic work, primarily benefitting students aiming to pursue a subsequent PhD or further career in academia. Yet, not all undergraduate students necessarily have this goal, with many considering other career paths in the Earth Sciences sector available. While the applied methodology of the case study was transferrable to other questions, the overall approach can easily be adapted for a different context.

From a practical and strategic point of view, the successful completion of the data collection activity resulting in a scientific publication provides a further tangible advantage. In times when funding for post-graduate positions is dwindling and the job market is increasingly competitive, having a publication on one’s CV is becoming essential (Berg, 2015). A co-authored publication resulting from a citizen science teaching project could therefore be beneficial for career progression however, not without the risk of exacerbating an already neoliberalised job market even further. This poses a major challenge for applicants from underrepresented and minority groups who face substantial hurdles in STEM subjects (Miller et al., 2019; Posselt et al., 2019). Extracurricular research, such as summer internships or fellowships, has been shown to be an effective tool to combat this attainment gap (Bruthers & Malays, 2020), but the lack of access to these can be another problem (Slovacek et al., 2011; Mogk, 2021). The citizen science teaching approach could partially mitigate this by providing students with research experience as part of the actual curriculum, eliminating the additional time and financial requirements often associated with external opportunities.

From a teaching perspective, the benefits of incorporating a citizen science approach in teaching are comparable to those of traditional citizen science projects. Data can be collected quickly and more effectively harnessing the efforts of multiple students. However, unlike the general public, undergraduate students already possess subject-specific knowledge. Furthermore, the required training can be highly interactive and specific, ensuring correctness and accuracy in data collection and with the added benefit of responding to the needs of individual students. A second, and not to be underestimated factor, is that multiple students collecting data alongside the expert teacher allows for the development and discussion of new ideas and feedback to improve the original project.

Despite these benefits, certain aspects warrant attention. As with traditional citizen science projects, there is a risk of exploiting cheap labour and issues with the recognition of knowledge creators and participants. For large-scale citizen science projects it is often impossible to include every single participant as a co-author (but see Sarna-Wojcicki et al., 2017). However, undergraduate courses in Earth Science disciplines are typically smaller making it feasible to manage co-ship more effectively (nevertheless, student numbers should be taken into account when designing similar projects as outlined below). Professional ethics and standards for co-authorships, such as the Vancouver Protocol, which attributes authorship to the acquisition, analysis, or interpretation of data, should be followed here.

Further concerns may be raised with regard to data quality and integrity (Riesch & Potter, 2014; Aceves-Bueno et al., 2017). However, the evaluation of public citizen science projects indicates that these concerns may be unwarranted (Resnik et al.,2014; Kosmala et al., 2016). This is in part due to the large amount of data collected redundantly to minimise erroneous entries or the application of software tools to identify outliers in the data (Kosmala et al., 2016). In a higher education setting, either similar approaches can be employed (see Considerations for study design below) or can be identified individually considering the smaller pool of participants.

### Considerations for study design

The citizen science approach presented here focuses on a case study in a palaeobiology context but is applicable more broadly to other themes and topics in Earth Sciences. However, some aspects need to be considered when adopting this approach (compare Kosmala et al., 2016).

### Data source

First and foremost, the data source needs to be considered carefully. The data to be collected has to be available relatively easily and not require complicated equipment or tools. This may rule out laboratory or field settings or data collection requiring high-spec computer equipment (although this may depend on the facilities and equipment available to teachers and students). In the presented example, data was collected from published literature sources that are readily available to students (although paywalls could present problems as access to scientific literature is not distributed equally globally (Boudry et al., 2019)). However, it is possible that data could also be collected from physical (e.g. fossil specimens, rock, ash, or soil samples, maps, etc.) or digital sources (e.g. 3D models, satellite images, geophysical data) if available in large enough quantities. The extent of the data collection will depend on the allocated time, which will dictate how many specimens or other sources can be analysed, the complexity of the measurements and analyses, and the sample size per student. Ideally, the data collection should be challenging enough to engage and motivate students throughout the course, but not so complex as to discourage participation.

### Ensuring data accuracy

Accuracy and consistency in the data collection are further points that have to be considered carefully during study design. It is likely that students will be unfamiliar with the specific details of the data collection and subsequent analysis, although they will be more knowledgeable about the overall context compared to public citizen science projects. A comprehensive introduction to the topic and thorough training in the required tasks are therefore paramount and should precede any data collection (Prysby & Oberhauser, 2004, Swanson et al., 2016). In the presented case study, the topical context and the methods were introduced in the first course session using an example of the process, which was simultaneously recorded as a video. This allowed students to revisit the example at any time. In addition, a concise handout describing all relevant data collection and analysis steps was created and provided for the students. While this procedure will form a solid basis for the data collection, it will not account for possible problems students may encounter. For example, the method may have to be adjusted or modified for specific samples (e.g. in the current example, eye socket position was not always lateral so some species required the measurements to be taken in dorsal view). Regular checks of the collected data will therefore be necessary. However, in contrast to traditional citizen science projects, students can ask questions and receive feedback on a continuous basis as part of the course. This will allow responding to problems and avoiding uncertainties during the data collection phase, which has been shown to improve data quality in other citizen science projects (Westphal et al., 2006).

A further measure to ensure the accuracy of the data is to implement a replication approach during the collection phase (Kosmala et al., 2016). In the current example, each sample specimen was assigned to two separate students, and comparing their data helped identify significant discrepancies. To avoid completely duplicating data sets for students and their assignments, the composition of the sample groups was varied; for example, the approximately 100 theropod dinosaurs in the sample were split into five groups of approximately 20 each first by taxonomic criteria and, for replication, by temporal distribution or randomisation. However, this approach may not always be possible if the number of students in the course (e.g. <10 students) does not permit duplicating the data collection tasks. In this case, it is advisable to either increase the sample size per student (time permitting) or to reduce the overall sample size. Conversely, with large course sizes (>150-200 students), data replication will be less of a problem but requires other measures to handle the increased data volume.

### Student engagement

Student engagement is a key consideration. Participants of traditional citizen science projects actively seek out projects and want to contribute to the data collection. While teachers might expect the same from undergraduate students, realities may differ from this perception, especially in the context of mandatory courses. However, this is not a problem unique to the introduced teaching approach but applies to any learning and teaching method in (higher) education and a plethora of studies have attempted to tackle this issue (Hattie, 2009; Fullan & Langworthy, 2013). Learning centered around case studies has been found to have a positive effect on student engagement (Prince, 2004; McMellon, 2013) and is underpinning the citizen science-style learning approach outlined here.

## Conclusions

Research-led/research-informed teaching has been at the center of countless pedagogical initiatives to improve teaching and learning in higher education. A citizen-science style approach to teaching embraces several key strategies at the heart of these pedagogical concepts, presenting a promising avenue for actively involving undergraduate students in research. The case study introduced here allowed students to develop a range of skills, including critical thinking, data collection and analysis, teamwork, and communication. The project was shown to enhance student engagement by providing meaningful, hands-on experiences that connect classroom learning to real-world applications with students often being more motivated when they understood that their work contributed to broader scientific endeavors. By integrating a case study in the context of an active research project into the curriculum, students were not only exposed to the practical aspects of research but also engaged in a meaningful investigation that contributed to the broader understanding of dinosaur biology and behavior. A citizen-science approach integrated into higher education curricula can therefore play a key role in the teaching and learning of Earth Sciences more generally and bridging the gap between research and teaching, transforming students from passive audiences to active participants in both learning and research activities.

## Acknowledgements

I would like to thank the year 1 undergraduate cohort 2022/23 of the palaeontology programme who engaged with the case study with open minds and full of motivation making the project a full success.

## Authors’ contributions

SL designed the study, carried out the analyses and wrote the manuscript. The author read and approved the final manuscript.

## Availability of data and materials

All data and material generated or analysed in this study are available in the publication

## Ethics approval and consent to participate

The study was carried out in accordance with the ethics guidelines of the University of Bristol.

## Consent for publication

Not applicable.

## References

Aceves-Bueno E, Adeleye AS, Feraud M, Huang Y, Tao M, Yang Y, Anderson SE. The accuracy of citizen science data: a quantitative review. Bull Ecol Soc Am. 2017;98(4):278–90.

Berg LD. Rethinking the PhD in the age of neoliberalization. GeoJournal. 2015;80(2):219–24.

Bonney R, Cooper CB, Dickinson J, Kelling S, Phillips T, Rosenberg KV, Shirk J. Citizen science: a developing tool for expanding science knowledge and scientific literacy. BioScience. 2009 Dec 1;59(11):977–84.

Boudry C, Alvarez-Muñoz P, Arencibia-Jorge R, Ayena D, Brouwer NJ, Chaudhuri Z, Chawner B, Epee E, Erraïs K, Fotouhi A, Gharaibeh AM. Worldwide inequality in access to full text scientific articles: the example of ophthalmology. PeerJ. 2019;7:e7850.

Brew A. Enhancing the quality of learning through research-led teaching. In Workshop presented at Annual Conference of the Higher Education Research and Development Society of Australasia: Quality Conversations, Perth, WA 2002 Jul.

Bruthers CB, Matyas ML. Undergraduates from underrepresented groups gain research skills and career aspirations through summer research fellowship. Adv Physiol Educ. 2020.

Carroll JM, Borge M, Xiao L, Ganoe CH. Realistic learning activity is not enough. In: 2008 Eighth IEEE International Conference on Advanced Learning Technologies, IEEE; 2008. p. 3–7.

Cohn JP. Citizen science: Can volunteers do real research?. BioScience. 2008 Mar 1;58(3):192–7.

Collins A. Cognitive Apprenticeship: Making Things Visible. Am Educ. 1991;15(3).

Committee on Higher Education. Higher Education Report of the Committee Appointed by the Prime Minister under the Chairmanship of Lord Robbins, 1961-1963. London: Cmnd 2154; 1963. p. 555–6.

Department for Education and Skills. The future for higher education. Norwich: The Stationery Office; 2003. Available at: http://www.dfes.gov.uk/highereducation/hestrategy/

Elton L. Research, Teaching and Scholarship in an expanding higher education system. High Educ Q. 1992;46(3):252–67.

Fullan M, Langworthy M. Towards a New End: New Pedagogies for Deep Learning. Collaborative Impact, Seattle, Washington, USA; 2013.

Griffiths R. Knowledge production and the research–teaching nexus: The case of the built environment disciplines. Studies in Higher education. 2004;29(6):709–26.

Healey M, Blumhof J, Thomas N. The Research-Teaching Nexus in Geography, Earth and Environmental Sciences (GEES). Planet. 2003;11(1):5–10.

Hattie JAC. Visible Learning. A synthesis of over 800 meta-analyses relating to achievement. London: Routledge Taylor & Francis Group; 2009.

Jenkins A, Blackman T, Lindsay R, Paton-Saltzberg R. Teaching and research: Student perspectives and policy implications. Studies in Higher education. 1998;23(2):127–41.

Kosmala M, Wiggins A, Swanson A, Simmons B. Assessing data quality in citizen science. Front Ecol Environ. 2016;14(10):551–60.

Kuh GD. What Student Affairs Professionals Need to Know About Student Engagement. J Coll Stud Dev. 2009;50(6):683–706.

Lautenschlager S, Aston RF, Baron JL, Boyd JR, Bridger HW, Carmona VE, Ducrey T, Eccles O, Gall M, Jones SA, Laker-McHugh H. Orbit size and estimated eye size in dinosaurs and other archosaurs and their implications for the evolution of visual capabilities. Journal of Vertebrate Paleontology. 2023 May 4:e2295518.

McMellon C. New advantages and insights into the living case teaching method: an exploratory study. J Acad Bus Econ. 2013;13(1):17–24.

Michelucci P, Dickinson JL. The power of crowds. Science. 2016 Jan 1;351(6268):32–3.

Miller CW, Zwickl BM, Posselt JR, Silvestrini RT, Hodapp T. Typical physics PhD admissions criteria limit access to underrepresented groups but fail to predict doctoral completion. Sci Adv. 2019;5(1):eaat7550.

Mogk DW. The intersection of geoethics and diversity in the geosciences. Geol Soc Lond Spec Publ. 2021;508(1):67–99.

Neumann R. The teaching-research nexus: Applying a framework to university students’ learning experiences. European Journal of Education. 1994;29(3):323–38.

Prince M. Does active learning work? A review of the research. J Eng Educ. 2004;93(3):223–31.

Posselt JR, Hernandez TE, Cochran GL, Miller CW. Metrics first, diversity later? Making the short list and getting admitted to physics PhD programs. J Women Minor Sci Eng. 2019;25(4).

Prysby MD, Oberhauser KS. Temporal and geographic variation in monarch densities: citizen scientists document monarch population patterns. In: Oberhauser KS, Solensky MJ, editors. Monarch butterfly biology & conservation. Ithaca, NY: Cornell University Press; 2004.

Resnik DB, Elliott KC, Miller AK. A framework for addressing ethical issues in citizen science. Environ Sci Policy. 2015;54:475–81.

Riesch H, Potter C. Citizen science as seen by scientists: Methodological, epistemological and ethical dimensions. Public Underst Sci. 2014;23(1):107–20.

Sarna-Wojcicki D, Perret M, Eitzel MV, Fortmann L. Where are the missing coauthors? Authorship practices in participatory research. Rural Sociol. 2017;82(4):713–46.

Slovacek SP, Whittinghill JC, Tucker S, Rath KA, Peterfreund AR, Kuehn GD, Reinke YG. Minority students severely underrepresented in science, technology, engineering, and math. J STEM Educ Innov Res. 2011;12(1).

Schapper J, Mayson SE. Research-led teaching: Moving from a fractured engagement to a marriage of convenience. Higher Education Research & Development. 2010 Dec 1;29(6):641–51.

Swanson A, Kosmala M, Lintott C, Packer C. A generalized approach for producing, quantifying, and validating citizen science data from wildlife images. Conserv Biol. 2016;30(3):520–31.

Powell MC, Colin M. Participatory paradoxes: facilitating citizen engagement in science and technology from the top-down? Bull Sci Technol Soc. 2009;29(4):325–42.

Vohland K, Weißpflug M, Pettibone L. Citizen Science and the Neoliberal Transformation of Science-an Ambivalent Relationship. Citizen Science: Theory & Practice. 2019;6(1).

Vohland K, Land-Zandstra A, Ceccaroni L, Lemmens R, Perelló J, Ponti M, Samson R, Wagenknecht K. Editorial: The Science of Citizen Science Evolves. 2021.

Westphal AJ, Von Korff J, Anderson DP, Alexander A, Betts B, Brownlee DE, et al. Stardust@ home: virtual microscope validation and first results. In: 37th Annual Lunar and Planetary Science Conference; 2006 Mar; p. 2225.

Zamorski B. Research-led teaching and learning in higher education: A case. Teaching in higher education. 2002;7(4):411–27.

